# Metabolic reorganization shapes the dynamics of a water kefir microbial community

**DOI:** 10.64898/2026.10.09.757983

**Authors:** Marine Chambeaudie, Delphine Mazé, Marie Cariou, Evelyne Duvernois-Berthet, Julien Mozziconacci, Ekaterina Tulaeva, Mathilde Courel, Emmanuelle Delagoutte, Fernando Lopes, Christophe Lavelle, Jean-Baptiste Boulé, Alexandra Joubert

**Affiliations:** Museum National d’Histoire Naturelle, CNRS UMR 7196, STRING, INSERM U1154, Sorbonne Université, Paris, France; Museum National d’Histoire Naturelle, CNRS UAR 2047, DOHNEE, Paris, France; Museum National d’Histoire Naturelle, CNRS UMR7221, PHYMA, Paris, France; Laboratoire Reproduction et Développement des Plantes, CNRS UMR 5667, École Normale Supérieure de Lyon, INRAE, Université Claude Bernard, Lyon, France

**Keywords:** water kefir, microbial community, metabolic reorganization, fermentation dynamics, carbon metabolism

## Abstract

Microbial fermentations emerge from the collective activity of interacting species whose relative abundances, metabolic functions, and environmental constraints continuously change over time. Water kefir provides a tractable model to investigate how multispecies communities coordinate their metabolic activities during fermentation. Here, we combined time-resolved community profiling, physicochemical measurements, metabolite quantification, and controlled co-culture assays to characterize the dynamics of a water kefir consortium comprising 19 detected microbial species. Our results reveal that water kefir fermentation is not adequately described as a simple succession of independently acting microbial groups. Instead, the community undergoes a reproducible reorganization of its collective metabolism around 20 h of fermentation. This transition marks a shift from an early regime characterized by the accumulation of soluble sugars and ethanol to a later regime dominated by enhanced sugar consumption and lactate production. Controlled interaction assays further showed that bacterial partners can alter carbon-use behaviour in *Saccharomyces cerevisiae*, highlighting the capacity of interspecies interactions to reshape metabolic phenotypes within the consortium. Together, these results support an ecological view of water kefir fermentation in which community-wide metabolic reorganization emerges from the interplay between microbial interactions, resource availability, and environmental change.

## 1 Introduction

Microbial communities are dynamic systems in which the fate of a resource rarely depends on the metabolism of a single species. Instead, substrates are transformed through networks of competition, facilitation, cross-feeding, inhibition, and environmental modification, so that the activity of one population continuously changes the ecological landscape experienced by the others. The resulting community function therefore cannot always be inferred from a simple inventory of the species present. It emerges from the temporal ordering of metabolic activities and from the feedbacks that couple resource availability, population structure, and physicochemical conditions (Harcombe *et al*. 2014, Ponomarova & Patil 2015, Gralka *et al*. 2020, Dal Bello *et al*. 2021). A central problem in microbial ecology is consequently to move from a taxonomic description of a consortium to a dynamical description of the transformations that organize it: which resources are consumed, which compounds are released, when the dominant metabolic processes change, and whether such changes occur progressively or through identifiable reorganizations of the community as a whole.

Water kefir, also referred to as sugary kefir or fruit kefir, is especially attractive in this respect because propagation of the microbial community and production of the fermented beverage are inseparable. A typical fermentation is initiated by transferring translucent gelatinous grains into an aqueous sucrose solution, commonly supplemented with dried or fresh fruit and sometimes lemon (Figure 1). During one to several days at room temperature, the inoculated medium becomes acidic, carbonated, weakly alcoholic, and enriched in microbial metabolites; the grains increase in mass and are subsequently recovered to initiate the next batch (Fiorda *et al*. 2017, Lynch *et al*. 2021, Zannini *et al*. 2023). The grain is therefore both an inoculum and a persistent physical habitat. Water kefir has recently been highlighted as a tractable system for integrative multi-omics and community metabolic analysis (Khan *et al*. 2026).

**Figure 1:**
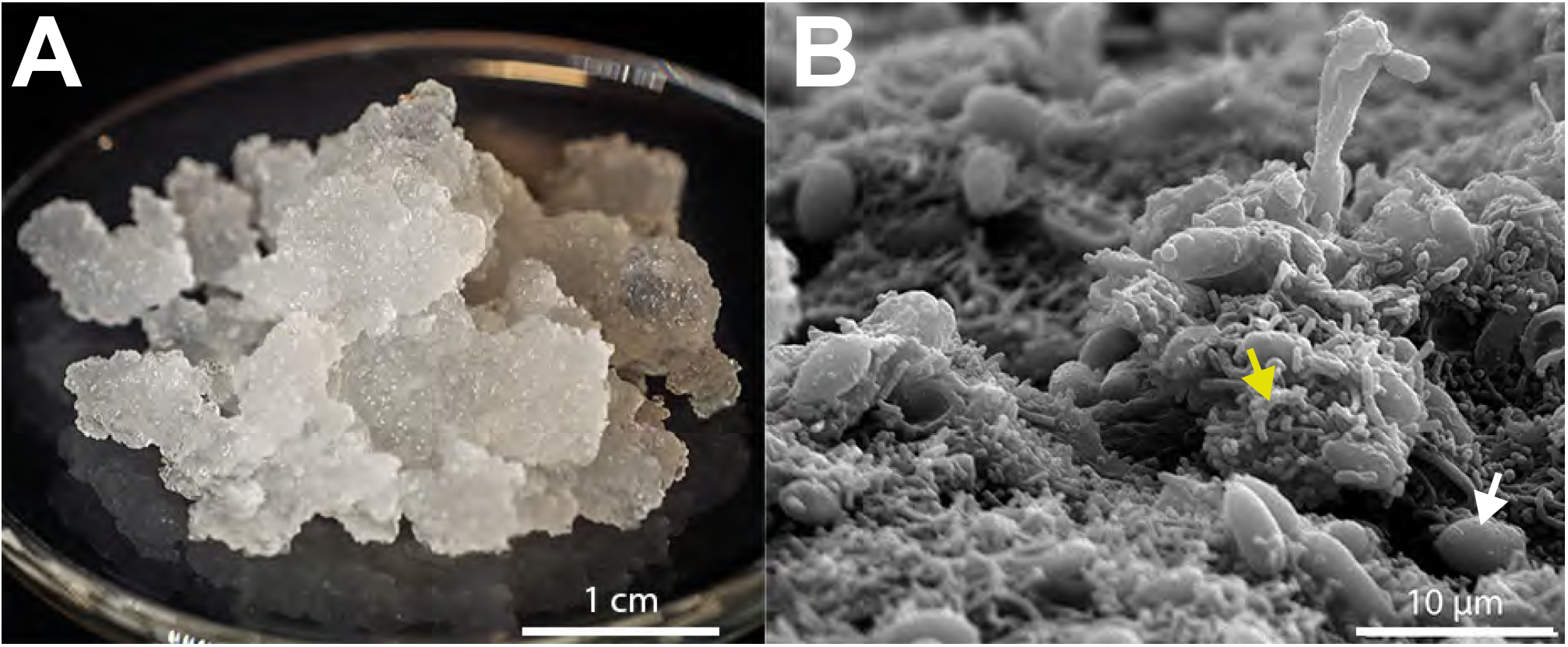
Water kefir grains and their microbial structure. **A**. Macroscopic appearance of water kefir grains. **B**. Scanning electron microscopy image of the grain surface showing yeast cells (white arrow) and bacterial cells (yellow arrow) embedded within the grain structure. Magnification: ×2500.

It is tempting to convert these known physiological capacities into a simple succession; yeasts first, lactic acid bacteria (LAB) second, acetic acid bacteria (AAB) last; but such a scheme is only a working model. In a mixed community, production and consumption occur concurrently, extracellular concentrations represent the net balance of several processes, and the physicochemical environment itself changes as a consequence of microbial activity. Measurements of metabolite concentrations must therefore be interpreted dynamically rather than as direct proxies for the activity of a single taxon. This is particularly important because most previous descriptions of water kefir have emphasized composition, endpoint chemistry, or individual metabolite trajectories (Laureys & De Vuyst 2014, Fiorda *et al*. 2017, Laureys & De Vuyst 2018, Verce *et al*. 2019, Patel *et al*. 2022, Zannini *et al*. 2023, Breselge *et al*. 2025b), while recent multi-omics studies have begun to resolve microbial succession and functional dynamics during water kefir fermentation (Arrieta-Echeverri *et al*. 2023, Breselge *et al*. 2025a, Garmendia *et al*. 2025); however, the existence, timing, and robustness of a community-wide metabolic transition remain less explicitly quantified.

These considerations motivate a shift in emphasis from microbial succession to metabolic reorganization. Rather than asking only which group dominates at each time point, one can ask how several measurable metabolites (produced or degraded) change dynamically during fermentation. If the rates of appearance and disappearance of multiple metabolites are considered jointly, a transition should manifest as a coordinated redistribution of apparent fluxes in multivariate space. Such an approach is conceptually close to viewing microbial communities as metabolically cohesive systems, in which ecosystem-level behaviour results from coupled transformations distributed across several populations (Pascual-Garćia *et al*. 2020, Machado *et al*. 2021). It also provides a way to distinguish a gradual drift from a reproducible change of regime and, *a posteriori*, to relate that transition to changes in oxygen, pH, substrate availability, community composition, or interspecies signalling.

Here, we use a representative domesticated water kefir consortium, referred to as MCF, to connect genomic composition, community dynamics, physicochemical evolution, metabolite kinetics, and interspecies effects within a single experimental framework. Time-resolved fermentations are subsequently used to follow microbial abundance together with pH, dissolved oxygen, sugars, ethanol, and organic metabolites. Rather than imposing an *a priori* number of fermentation phases, we analyse the joint dynamics of the measured carbon compounds and test whether the resulting apparent fluxes identify a reproducible transition in community metabolism. Finally, controlled interaction assays with *S. cerevisiae* examine whether bacterial partners can modify yeast carbon-use phenotypes, thereby providing an experimental link between interspecies interactions and metabolic plasticity.

## 2 Materials and Methods

### Water kefir consortium and controlled fermentations

The consortium selected for the study originated from a household water kefir culture maintained for more than ten years in Paris, France, before transfer to the laboratory. This consortium is hereafter referred to as MCF. Following transfer, MCF grains were propagated continuously under standardized laboratory conditions.

Unless otherwise indicated, MCF grains were inoculated at 40 g.L^−1^ into a standardized fig-lemon medium. The medium consisted of autoclaved Paris tap water (Tables S1 and S2), supplemented with one dried organically produced fig per litre, 0.5 mL.L^−1^ organic lemon juice and 40 g.L^−1^ sucrose. Routine batch fermentations were performed at 25°C without agitation. Detailed procedures for the biodiversity survey, cultivation, scanning electron microscopy, microbial isolation, DNA extraction, sequencing, genome reconstruction and Hi-C analysis are provided in the Supplementary Methods.

### Physicochemical, community and metabolite kinetics

The physicochemical evolution of the fermentation was monitored for 96 h in a bioreactor set-up at 25°C without mechanical agitation. Calibrated probes continuously recorded pH, dissolved oxygen, oxido–reduction potential (ORP) and temperature. Details on the bioreactor set–up and probes used in this study are given in Supplementary Methods.

Microbial dynamics were examined independently in the grain-associated and liquid-phase communities. Experiments were performed in two sequential biological replicates for grain kinetics and in three sequential biological replicates for beverage kinetics at 10, 30, 48, and 72 h. Beverage pellets obtained after centrifugation, as well as the washed grains, were processed as described in Supplementary Methods for DNA extraction and long-read sequencing. To obtain strain-resolved estimates, all contigs longer than 1 kb from the 19 reconstructed MCF genomes were concatenated to produce a synthetic reference metagenome. Long reads from each kinetic sample were mapped against the complete synthetic metagenome using Minimap2 (Li 2018, Supplementary Methods). These values were interpreted as relative community representation rather than as direct measurements of absolute cell number.

Fermentation metabolites were quantified at 0, 5, 10, 15, 20, 24, 30, 35, 48 and 72 h. Sucrose, D-glucose, D-fructose, glycerol, gluconate, mannitol, lactate and acetate were quantified enzymatically using commercial assay kits according to the manufacturers’ recommendations (Megazyme#K-SUFRG, #KGATE, #K-MANOL, #K-DLATE and Sigma-Aldrich #MAK086). Samples were processed using a Uvikon XL Secoman spectrophotometer, except for gluconate and acetate, for which the protocol was adapted for measurement in microplate in a Perkin Elmer Wallac 1420 victor2 Microplate Reader set to the recommended wavelength. Ethanol was quantified in parallel over the same kinetic series using an inline PASPORT Ethanol Sensor (PASCO). Unless otherwise indicated, metabolite measurements were performed in triplicate and are presented as mean values with the corresponding experimental dispersion.

### Bacteria–yeast interaction assays

The effect of bacterial partners on carbon-source utilization by MCF yeasts was examined using an agar-plate assay adapted from experimental approaches developed for the glucose-associated repression (GAR) assay in *Saccharomyces cerevisiae* (Brown & Lindquist 2009, Jarosz *et al*. 2014, Watanabe & Takagi 2019). Full culture conditions and plate compositions are provided in the Supplementary Methods.

### Apparent extracellular carbon fluxes and metabolic-state representation

To examine the collective dynamics of extracellular carbon redistribution independently of the absolute concentration scale of each compound, the kinetic measurements were recast as carbon-containing pools. This analysis was restricted to the nine metabolites sharing a strict common experimental time grid without interpolation: sucrose, glucose, fructose, ethanol, acetate, lactate, mannitol, glycerol and gluconate. For metabolite *i*, its measured concentration *c*_*i*_ (*t*) in g.L^−1^ was converted into an equivalent carbon concentration *C*_*i*_ (*t*), expressed in mmol-C.L^−1^, according to,

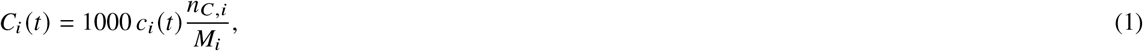

where *n*_*C,i*_ is the number of carbon atoms contained in one molecule of metabolite *i* and *M*_*i*_ is its molar mass in g.mol^−1^. This conversion is strictly dimensional and does not impose a biochemical or stoichiometric model. In biological terms, it simply places all measured compounds on a common carbon scale before their temporal changes are compared.

The net apparent flux of each measured carbon pool between two consecutive observations was estimated by finite differences,

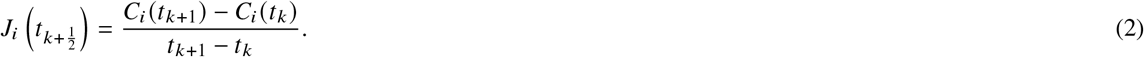

Accordingly, *J*_*i*_ > 0 denotes net extracellular accumulation of compound *i* over the corresponding interval, whereas *J*_*i*_ < 0 denotes net depletion. These quantities are referred to throughout as apparent extracellular fluxes; they do not represent intracellular metabolic fluxes in the strict biochemical sense.

Because the amplitudes of the measured pools differ substantially, each metabolite flux was standardized over the complete kinetic series,

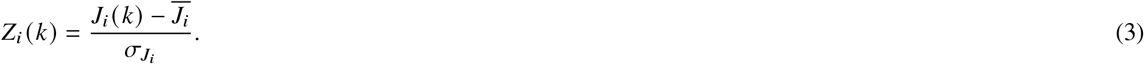

Each interval was therefore represented by a dimensionless nine-component vector **Z**(*k*). Standardization gives comparable weight to metabolites with very different absolute concentration scales; the vector therefore describes the pattern of extracellular carbon accumulation and depletion during each time interval rather than the abundance of one dominant compound.

### Objective detection of a metabolic transition

Two independent criteria were used to identify the dominant transition in the multivariate apparent-flux trajectory without imposing an *a priori* transition time. First, the magnitude of the local change between successive metabolic states was quantified as,

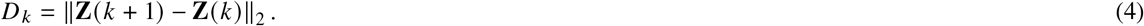

The sampling boundary associated with the largest value of *D* _*k*_ defines the strongest local reorientation of the multivariate apparent-flux vector. Put more simply, this criterion asks when the combined pattern of production and depletion changes most abruptly from one sampled interval to the next.

Second, a global single-change-point model was evaluated (Page 1954, Truong *et al*. 2020). For each admissible candidate boundary *τ*, the standardized flux vectors were separated into a pre-transition set *P*(*τ*) and a post-transition set *Q*(*τ*). The total within-regime dispersion was calculated as,

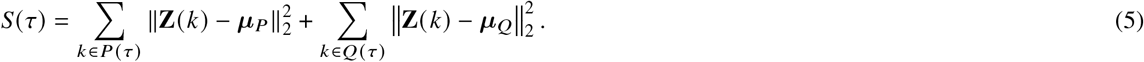

The preferred change point was defined as the candidate boundary minimizing *S*(τ). Thus, the local and global criteria interrogate different properties of the trajectory: the former identifies its largest adjacent change, whereas the latter identifies the partition producing the most internally coherent two-regime representation. Biologically, the second criterion asks where the fermentation trajectory can be divided into two internally most similar metabolic regimes.

For each metabolite, its contribution to the transition was summarized by the difference between its mean standardized apparent flux after and before the optimal boundary,

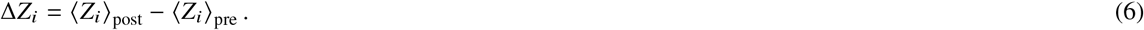

The resulting Δ*Z*_*i*_ values are dimensionless effect-size descriptors and were not interpreted as absolute biochemical fluxes. Robustness was evaluated by leave-one-metabolite-out analysis and by Monte-Carlo propagation of experimental uncertainty; full details are provided in the Supplementary Methods.

## 3 Results

### The MCF consortium lies within the diversity of water kefir microbial communities

More than one hundred water kefir grain samples were collected, primarily from private growers in France, and a subset of seventeen independent grains was subjected to an initial shotgun metagenomic survey. Short reads were mapped against a synthetic reference database comprising 71 bacterial and yeast genomes previously associated with water kefir. The seventeen grains displayed substantial quantitative variation while retaining a recurrent taxonomic structure (Figure 2). Several grains additionally contained Bifidobacterium species, *Zymomonas mobilis*, or both, taxa that have previously been detected in water kefir ecosystems (Patel *et al*. 2022, Breselge *et al*. 2025b). In the *Z. mobilis*-positive samples, this organism could represent a major fraction of the mapped signal, reaching approximately 80% of normalized genome coverage in the most extreme cases. The MCF grain lacked detectable *Z. mobilis* and occupied the range defined by the other *Zymomonas*-free communities. Its composition was therefore not exceptional within our survey, while its long maintenance history and reproducible laboratory propagation made it suitable for detailed experimental characterization. We consequently selected MCF as a model consortium for the remainder of the study.

**Figure 2:**
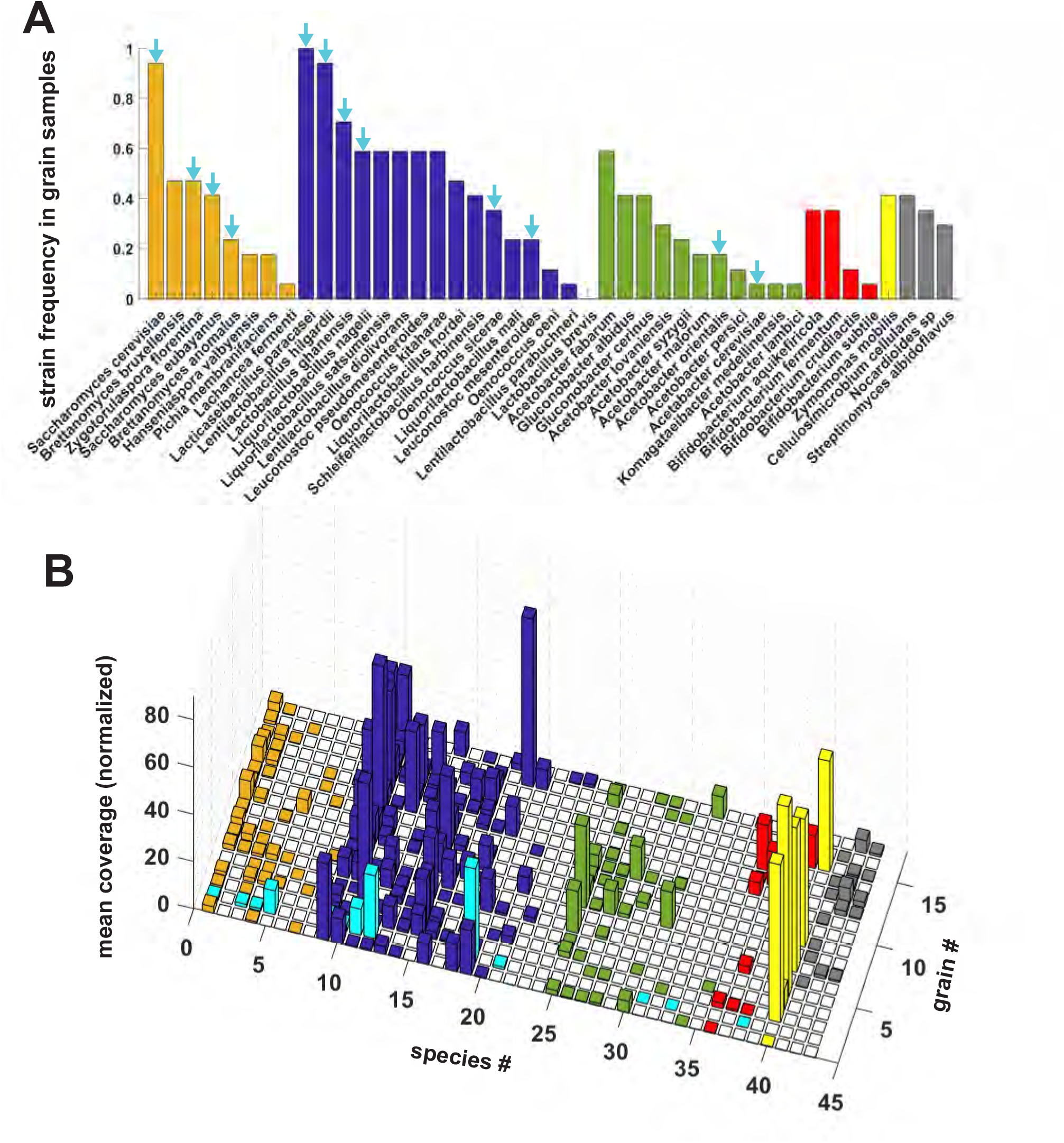
Microbial diversity among water kefir grain samples. Genomic DNA from 17 independently maintained water kefir grains was sequenced using Illumina technology and mapped against a reference database containing 71 genomes representative of microorganisms previously reported in water kefir. Only the species detected in the water grains kefir are represented here. **A**. Frequency of detection of individual taxa among the 17 grains. The cyan arrows indicate the species detected in the MCF consortium. **B**. Normalised genome coverage for each reference taxon in each grain sample. The cyan-highlighted sample corresponds to the MCF consortium.

### A near-complete genomic catalogue resolves 19 members of the MCF consortium

We next combined culture-dependent isolation, long- and short-read sequencing, community metagenomics and Hi-C information to resolve the microbial composition of the MCF consortium at strain level. Eighteen microorganisms could be isolated and maintained independently (Supplementary Figure S1). A nineteenth organism, *Oenococcus sicerae*, was consistently detected in community sequencing data but could not be maintained as an isolated culture under the conditions used here. Its genome was therefore reconstructed directly from metagenomic reads. The final assemblies were of high quality, with most bacterial genomes assembling into single circular chromosomes, along with potential plasmids (Supplementary Table S4), while yeast genomes assembled into large scaffolds. A metaHiC analysis was performed on the liquid phase after 30h to validate our consortium composition and improve genomes assemblies especially for the yeast species (Supplementary Figure S2-S5).

The resulting catalogue comprises 19 species: four yeasts, nine LAB and six AAB (Table 1). Together, cultivation and culture-independent reconstruction yielded a genomic reference encompassing the dominant detectable members of the MCF consortium and provided the strain-specific reference required for the subsequent time-resolved metagenomic analysis.

**Table 1:**
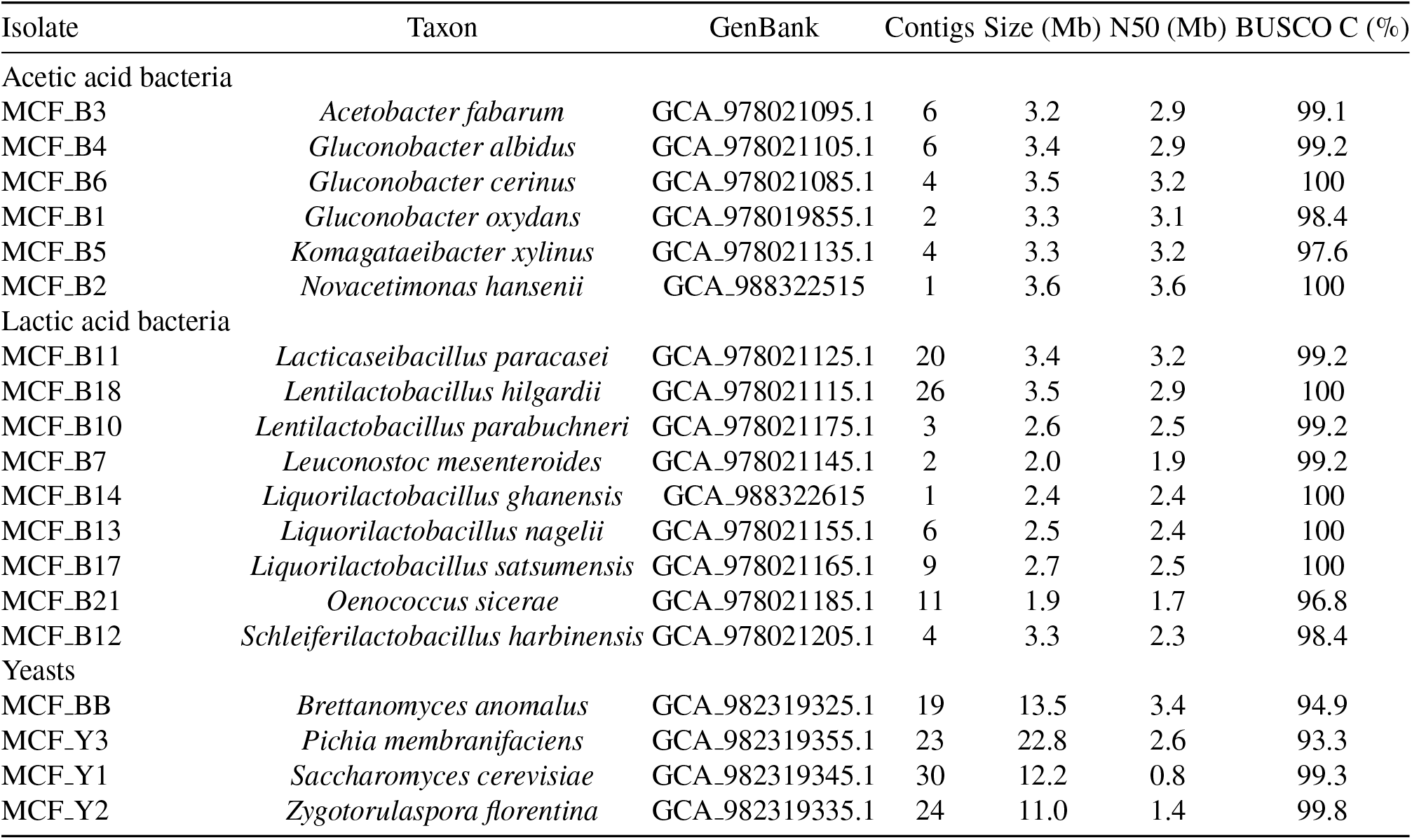
Genome assemblies obtained for the 19 microbial members of the MCF consortium. Genome completeness corresponds to the complete BUSCO fraction obtained with the appropriate lineage dataset (ENA PRJEB105405).

Interestingly, the combination of both metagenomic analysis and culture-dependent isolation of species present in MCF grains allowed us to refine the composition of the consortium. Owing to limitations of the metagenomic approach, 9/19 species were correctly identified by metagenomic analysis, while culture dependent approach allowed to retrieve 10 additional species poorly represented in the grain.

### Fermentation rapidly establishes an acidic and strongly reduced environment

Continuous measurements of physicochemical parameters revealed a highly reproducible physicochemical trajectory (Figure 3). pH progressively decreased from approximately 4.1 at inoculation to approximately 3.4 during the late fermentation period. Dissolved oxygen showed the most rapid environmental change. The medium initially contained measurable oxygen but the pO_2_ signal collapsed within the first hours and subsequently remained close to the detection limit. The combination of persistent oxygen depletion, acidification and decreasing ORP therefore shows that the fermentation rapidly evolves toward a reduced environment and remains strongly oxygen limited thereafter

### Community composition is comparatively stable in the grain but reorganizes in the beverage

We next examined how the physicochemical evolution of the fermentation was accompanied by changes in community composition. Strain-resolved mapping revealed strikingly different dynamics in the grain and in the surrounding liquid phase (Figure 4).

**Figure 3:**
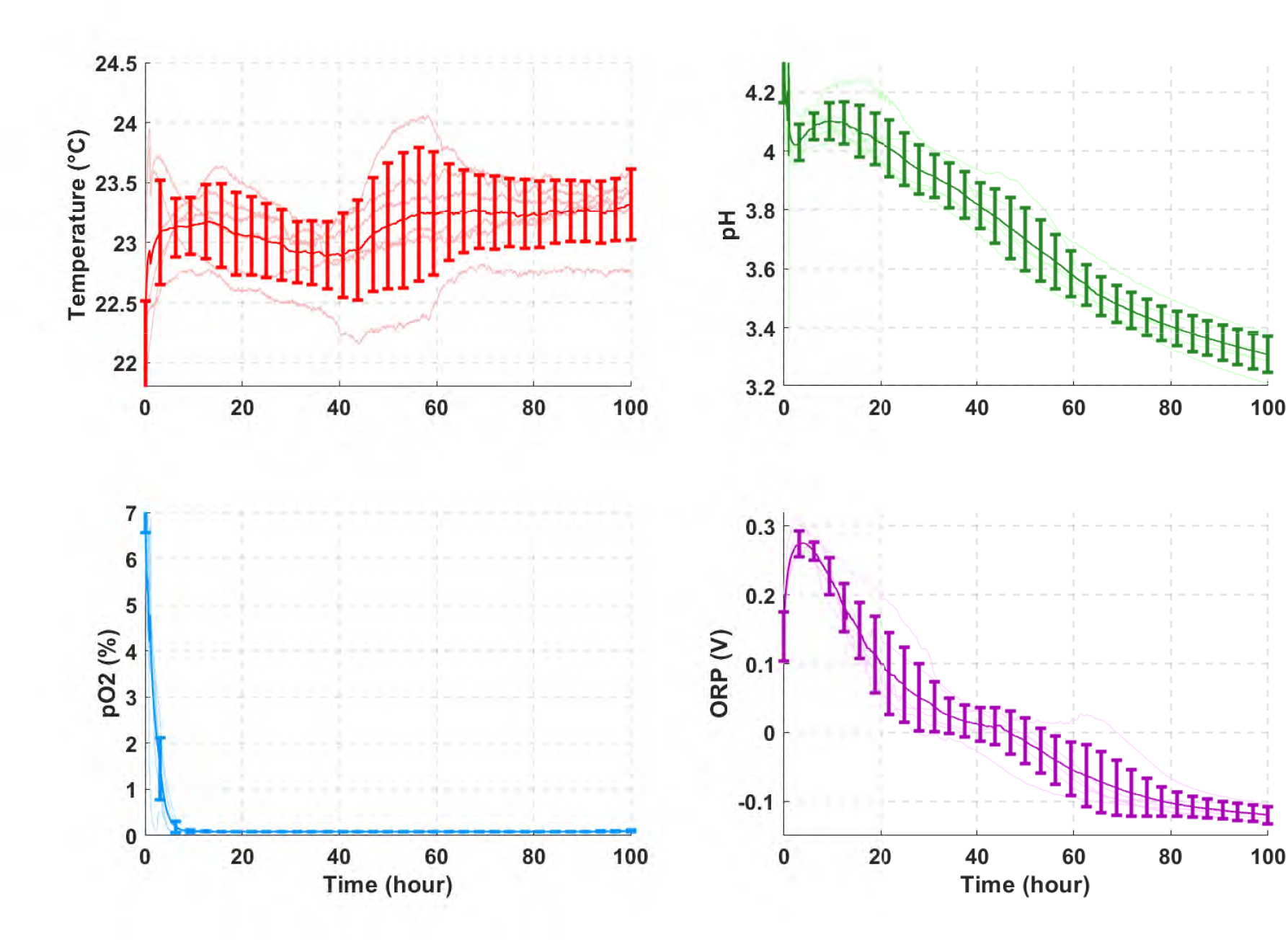
Physicochemical evolution of MCF water kefir fermentation. Temperature (°C), pH, dissolved oxygen (pO_2_) and Oxido-reduction potential (ORP, in V) were continuously monitored during fermentation in the bioreactor. Dissolved oxygen was rapidly depleted, approaching detection limit, while a steady decrease in pH and RedOx potential continued throughout the fermentation..

**Figure 4:**
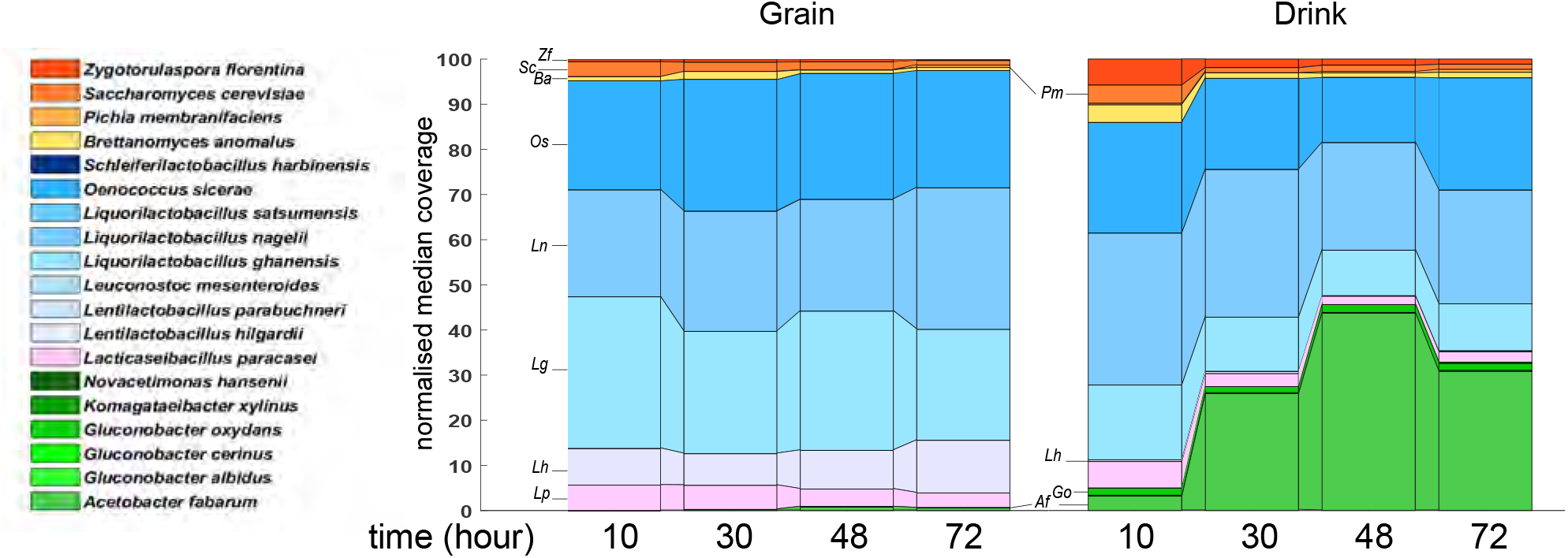
Temporal dynamics of microbial community composition in the MCF grain and fermented beverage. The grain-associated community remains comparatively stable, whereas the liquid phase pronounced reorganization over the course of fermentation. Values represent normalised median coverage distributions and not absolute cell abundances.

Within the grain, the community was dominated by LAB throughout the experiment. More importantly, the relative distribution of the dominant grain-associated taxa changed only modestly between 10 and 72 h. Thus, the grain retained a comparatively stable taxonomic structure during a single fermentation cycle despite the pronounced environmental changes occurring in the surrounding medium.

The beverage displayed a substantially more dynamic profile. At 10 h, yeasts including *S. cerevisiae, B. anomalus* and *Z. florentina* contributed appreciably to the mapped community. By 30 h, their relative representation had decreased while several LAB, including *L. paracasei, L. hilgardii, L. ghanensis, L. nagelii* and *O. sicerae*, became more prominent. A particularly pronounced enrichment was observed for *A. fabarum*. This organism represented only a minor fraction at 10 h but became one of the dominant taxa in the liquid phase between 30 and 48 h.

These values correspond to relative sequence representation and should not be interpreted as direct absolute cell counts. Nevertheless, they demonstrate that the taxonomic composition of the liquid phase is reorganized during fermentation whereas the grain-associated population remains comparatively stable.

### Metabolite kinetics reveal a progressive redistribution of carbon resources

The metabolic evolution of the same fermentation showed that changes in community composition were accompanied by a coordinated redistribution of extracellular carbon compounds (Figure 5). Sucrose, initially present at approximately 40 g.L^−1^, progressively disappeared and was almost completely exhausted by 72 h. In contrast, glucose and fructose accumulated during the first part of fermentation, consistent with sucrose hydrolysis, and reached their highest concentrations around 20 h. Their trajectories subsequently reversed, with both sugars progressively depleted during the remainder of the experiment. Most microorganisms present in the kefir consortium were able to grow on multiple carbon sources (glucose, fructose, sucrose, glycerol), albeit with varying efficiency (Supplementary Figure S6 and S7), suggesting for example that bacterial growth in the consortium does not solely depend on sucrose inversion by yeasts. Most bacteria also express dextransucrases, allowing matrix building from sucrose (Table S5).

**Figure 5:**
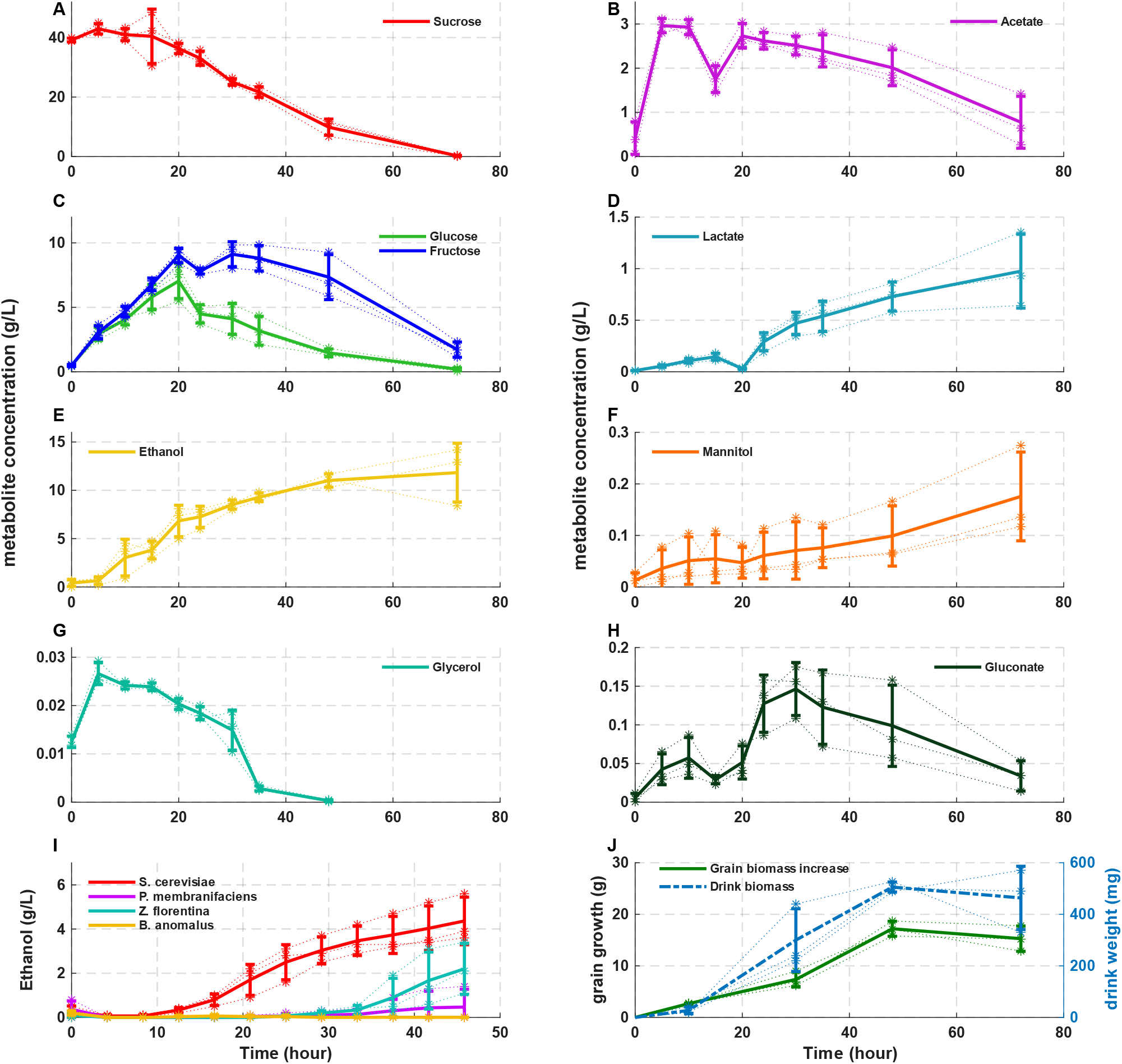
Temporal dynamics of substrates, fermentation products and biomass during MCF water kefir fermentation. Sucrose, acetate, glucose, fructose, lactate, ethanol, mannitol, glycerol and gluconate were followed through fermentation; panels also show ethanol accumulation by isolated yeasts and grain and beverage biomass. Error bars represent replicate dispersion.

Lactate followed an opposite trajectory. Its concentration remained low during the first part of the experiment but increased markedly after approximately 20 h and continued to accumulate thereafter, reaching close to 1 g.L^−1^ by 72 h. Mannitol also progressively accumulated, although at substantially lower concentrations. Gluconate increased during the intermediate fermentation period to approximately 0.15 g.L^−1^ and subsequently declined.

These trajectories argue against a simple interpretation in which each metabolite is associated with a single temporally isolated microbial group. In particular, ethanol continues to accumulate despite enrichment of AAB, while acetate decreases during the late fermentation period. The strongly oxygen-limited environment documented independently by the pO_2_ measurements provides a plausible physicochemical constraint on extensive oxidative conversion of ethanol to acetate (De Roos & De Vuyst 2018, da Silva *et al*. 2022). The MCF fermentation therefore does not undergo a classical late acetic fermentation under the conditions used here.

### A community-wide metabolic reorganization occurs around 20 h

Visual inspection of the individual metabolite trajectories suggested that several variables changed behaviour within a relatively narrow time interval around 20 h. In particular, glucose and fructose ceased to accumulate, lactate entered a sustained accumulation regime, and several lower-abundance metabolites showed concomitant changes in their rates of appearance or disappearance. We therefore asked whether this apparent transition could be identified objectively from the joint dynamics of the measured extracellular carbon pools.

The analysis was restricted to the nine compounds for which a strict common sampling grid was available between 0 and 48 h: sucrose, glucose, fructose, ethanol, acetate, lactate, mannitol, glycerol and gluconate. Concentrations were first converted to mmol-C.L^−1^ and finite differences were used to define apparent extracellular carbon fluxes (see Material and Methods). Because the absolute amplitudes of the different metabolite pools vary substantially, each flux was standardized before multivariate analysis.

Figure 6 illustrates the marked reorganization of the metabolic regime around 20 h. The multivariate metabolic-state score (Figure 6A) changes abruptly between the early and late phases of fermentation, with the transition occurring close to 20 h despite the propagated experimental uncertainty. The largest metabolic process change is an increase in overall sugar uptake (Δ*Z* = 1.49), calculated from the mean standardized changes in sucrose, glucose, and fructose consumption. This is accompanied by a strong increase in lactate production (Δ*Z* = 1.21). Glycerol uptake also increases substantially (Δ*Z* = 0.91), while ethanol production decreases by a comparable standardized magnitude (Δ*Z* = 0.91). Acetate production is likewise reduced (Δ*Z* = 0.64). In contrast, changes in gluconate and mannitol production are more modest (Δ*Z* = 0.20 and 0.16, respectively) (Figure 6B). Thus, the transition at approximately 20 h is characterized primarily by enhanced carbohydrate consumption and lactate production, together with reduced ethanol and acetate production, rather than by a uniform increase or decrease of all measured metabolic outputs. The timing of this transition is highly robust: 20 h is selected in 90.3% of the Monte-Carlo realizations, and all 9 leave-one-metabolite-out analyses independently retain the same transition time (Figure 6C).

**Figure 6:**
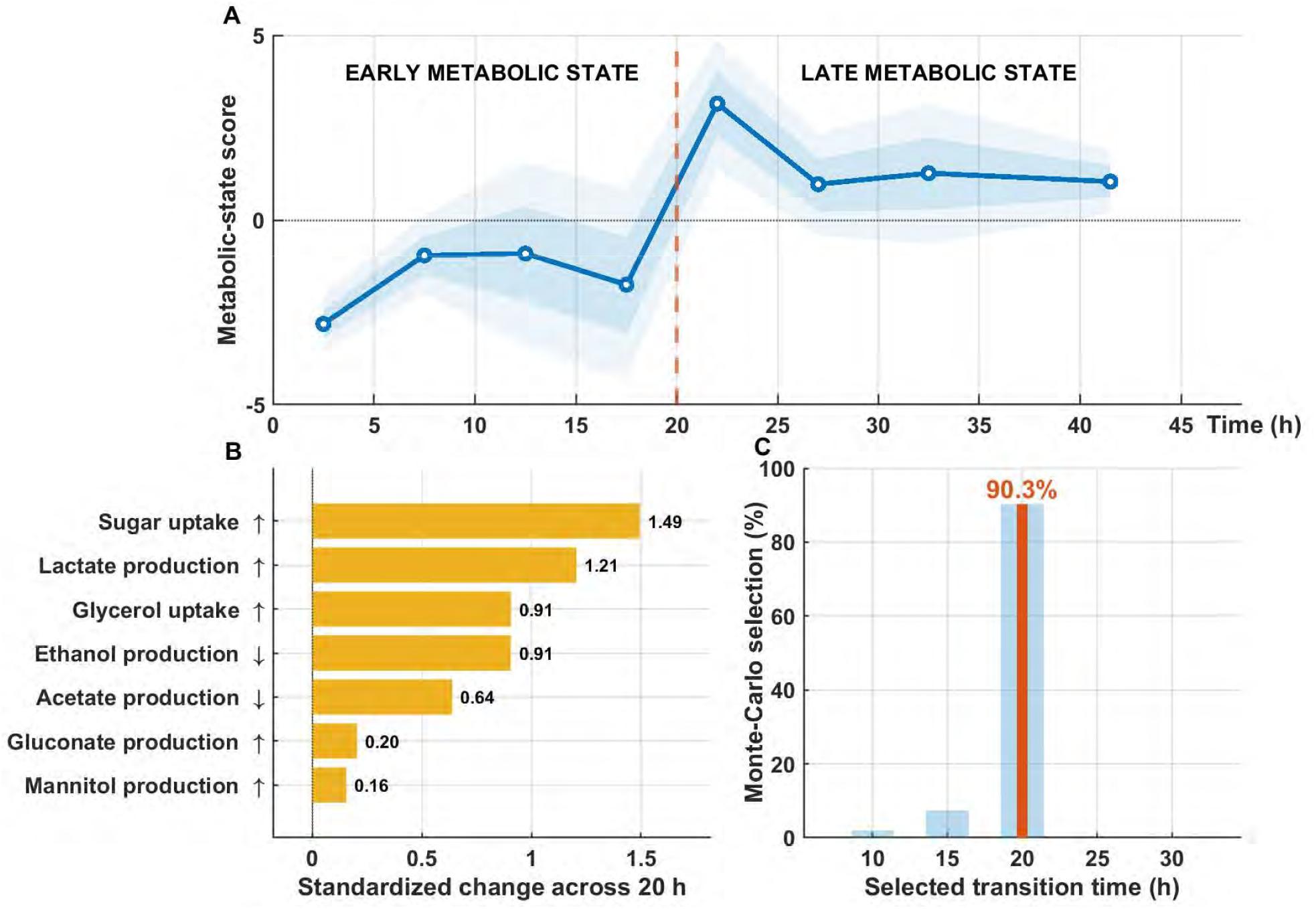
A dominant metabolic transition around 20 h reorganizes water kefir fermentation. **A**. Multivariate metabolic-state score obtained from the standardized apparent carbon fluxes. The dashed vertical line indicates the objectively selected transition time. Shaded regions represent the 68% and 95% uncertainty intervals propagated from the experimental concentration uncertainties. **B**. Main metabolic changes across the transition. Sugar uptake represents the mean standardized change in sucrose, glucose, and fructose consumption. Values correspond to standardized changes between the late and early metabolic regimes. For clarity, the signs in panel **B** were oriented according to the biological process indicated by each label, so that a positive bar always denotes a strengthening of the stated process after the transition; for example, increased uptake corresponds to a more negative concentration-derived flux, whereas decreased ethanol production is displayed as a positive magnitude of reduction. The transition is dominated by increased sugar uptake and lactate production, accompanied by increased glycerol uptake and decreased ethanol and acetate production, whereas gluconate and mannitol production show smaller changes. **C**. Robustness of the inferred transition time. Monte-Carlo resampling selects a transition at 20 h in 90.3% of realizations, while all 9 leave-one-metabolite-out analyses recover the same transition time.

The combined analyses therefore support a reproducible reorganization of the measured extracellular metabolic dynamics around 20 h. Before this boundary, hydrolysis and production of soluble sugars coexist with their consumption and ethanol accumulation. After the boundary, all three major soluble sugars shift toward stronger net depletion while lactate accumulation becomes substantially enhanced. Within the temporal resolution of the experiment, the data consequently support two distinguishable apparent-flux regimes separated by a dominant transition more clearly than they support three equally defined sequential metabolic phases.

Importantly, this result does not constitute a closed carbon balance and does not identify intracellular pathway fluxes. Nor does it, by itself, assign the transition to a particular microorganism. Rather, it establishes an objective temporal landmark at which multiple chemically distinct extracellular pools change their dynamics coherently.

### Bacterial partners alter carbon-use behaviour in *Saccharomyces cerevisiae*

The coordinated metabolic reorganization prompted us to ask whether interactions among consortium members could directly alter carbon utilization by one of the dominant yeasts. We focused on *S. cerevisiae* because previous studies have shown that bacterial interactions can modify glucose-associated repression and permit growth on alternative carbon sources under conditions in which glucose-related signalling would otherwise restrict their utilization (Brown & Lindquist 2009, Jarosz *et al*. 2014, Watanabe & Takagi 2019).

Co-culture with AAB altered this phenotype in a strain-dependent manner. As shown in Figure 7, *Gluconobacter cerinus* promoted detectable *S. cerevisiae* growth on GGM in its vicinity, whereas *Acetobacter fabarum* produced little or no comparable stimulation under the same conditions. Thus, the response was not a generic consequence of the presence of an AAB cell population but depended on the bacterial partner. All the other *S. cerevisiae* - bacteria combinations are shown in Supplementary Figures S8 and S9. These results demonstrate that bacterial partners present within the MCF consortium can modify the carbon-use behaviour of *S. cerevisiae*. They do not establish the molecular mechanism underlying this response. We tested if this phenotype could be transmitted transgenerationally (Supplementary Figure S10). Interestingly, even though the initial induction from LAB was lower than the induction from AAB, the reminiscent phenotype was higher in LAB induced growth and coherent with former observations. These results highlight the complex metabolic interplay between yeast and bacteria in Kefir consortia.

**Figure 7:**
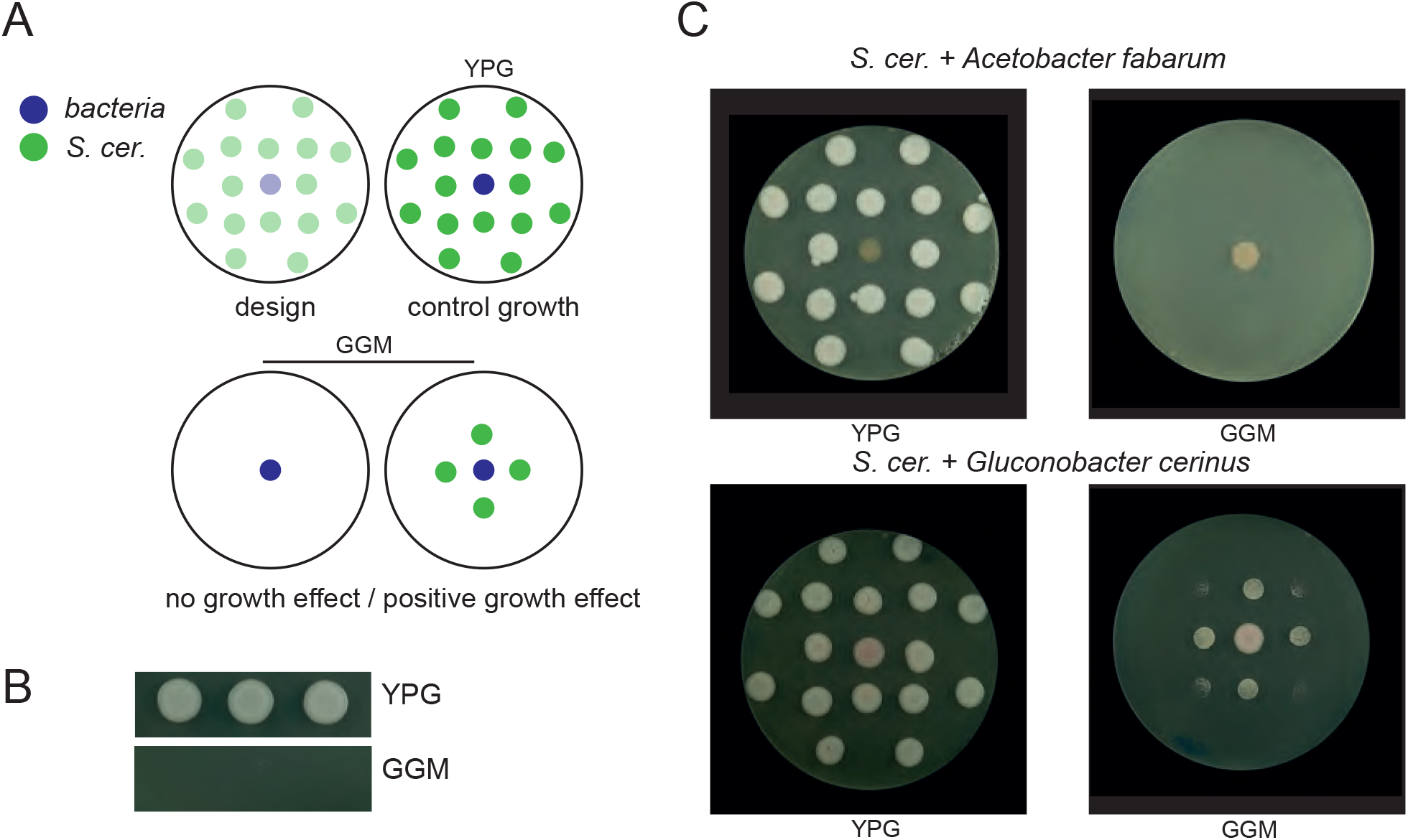
Acetic acid bacteria modify carbon-use behaviour of the MCF *Saccharomyces cerevisiae* isolate. **A**. Principle of the interaction assay. **B**. Basal growth on YPG and GGM. **C**. Representative interactions with *Acetobacter fabarum* and *Gluconobacter cerinus*.

## 4 Discussion

### Metabolic reorganization rather than simple succession

The observed dynamics require reconsideration of a common conceptual representation of water kefir fermentation as a sequence of yeast fermentation, lactic fermentation and subsequent acetic oxidation. Such a sequence is physiologically plausible because the community contains organisms capable of each of these processes. The presence of these capacities, however, does not establish that they occur as three temporally separated phases.

The distinction between compatibility and demonstration is important because the MCF community contains extensive functional redundancy. Several organisms can use the same carbon sources, and the same metabolite may be produced or consumed by different taxa. Such redundancy can make community-level function more stable than species abundance, because losses or fluctuations in one population may be partly compensated by another (Louca *et al*. 2018). This is a potential explanation for the coexistence of substantial taxonomic variability among independently maintained kefir grains with broadly conserved fermentation properties (Laureys & De Vuyst 2018, Zannini *et al*. 2023, Breselge *et al*. 2025b). It also provides a mechanistic basis for treating metabolic organization, rather than individual taxon abundance, as the more informative level at which to compare water kefir communities.

### Interactions modify carbon-use behaviour

Direct metabolic interactions between yeasts and LAB have previously been demonstrated in water kefir co-cultures, including growth effects associated with bacterial acidification and yeast-derived nutrients (Stadie *et al*. 2013). The interaction assays add a second dimension to this ecological coupling. They show that bacterial partners can alter the carbon-use phenotype of the MCF *S. cerevisiae* strain under restrictive glycerol-glucosamine conditions. The effect varies between the bacterial partners tested, indicating that the response depends on the identity of the interacting strain. This observation is particularly interesting because it shows that microbial interactions can directly modify carbon-use behaviour in another member of the community.

We do not claim that the 20-h transition is caused by induction of the yeast phenotype measured on plates. The temporal and mechanistic evidence required for such a causal inference is not available. Rather, the two results demonstrate complementary properties of the system: the consortium undergoes a collective change in extracellular carbon transformation, and its members are capable of modifying one another’s carbon-use behaviour. Whether these phenomena are mechanistically connected is a directly testable question.

### A transition-centered view of fermentation

Taken together, the results suggest a different conceptual representation of water kefir fermentation. In a purely succession-based model, community function is described primarily as the ordered replacement of organisms with different metabolic capacities. In the transition-centered model supported here, taxonomic composition, substrate availability and environmental conditions evolve continuously, but the collective extracellular metabolism occupies distinguishable regimes separated by a relatively sharp reorganization. Resource-dependent transitions between distinguishable microbial community states have also been observed experimentally in synthetic consortia, notably along gradients of total carbon availability (Bischofberger *et al*. 2026), although the mechanisms and observables considered here are different.

The two descriptions are not mutually exclusive. Microbial succession can contribute to metabolic transitions, and metabolic changes can in turn alter which populations are favoured. The important point is that neither level is sufficient alone. The community trajectory emerges from feedback between population dynamics and resource transformation.

The precision of the 20-h transition should nevertheless be interpreted in relation to the temporal resolution of the experiment. The analysis identifies the sampling boundary around 20 h as the dominant change point among the available observations; it does not imply that the biological system switches instantaneously at exactly 20.0 h. A more appropriate interpretation is that the fermentation crosses a relatively narrow transition region centred on this period. Detailed limitations and direct experimental extensions are provided in the Supplementary Note.

### A microbial biodiversity reservoir that promotes ecological antifragility

The ecological organization of the consortium is spatially differentiated. The grain retains a comparatively stable LAB-dominated community over the fermentation cycle, whereas the surrounding beverage undergoes substantially greater taxonomic redistribution. The grain thus appears to behave as a persistent microbial reservoir from which a more dynamic planktonic community develops. Such compartmentalization may be central to the long-term propagation of water kefir, because it allows substantial short-term ecological variation without requiring complete reconstruction of the consortium at every fermentation cycle. The interaction experiments further show that this community is not reducible to a set of organisms competing independently for the same substrates. Bacterial partners can modify carbon-use behaviour in *Saccharomyces cerevisiae*.

These observations lead to a more general view of fermented microbial communities. Stability need not imply immobility; microbial community stability is commonly distinguished in terms of properties such as resistance and resilience rather than strict compositional invariance (Shade *et al*. 2012). The MCF consortium remains functionally coherent while undergoing pronounced changes in its chemical environment, resource availability and liquid-phase population structure. What persists is not a fixed instantaneous state but the capacity to traverse a reproducible dynamical trajectory. In this sense, the relevant unit of stability may be the trajectory itself rather than any particular microbial composition measured at a single time point. This distinction invites comparison with the concept of antifragility introduced by Taleb (2012). Antifragility is stronger than robustness: a robust system withstands perturbation, whereas an antifragile system derives some functional advantage from variability, stress or repeated disturbance. Water kefir is propagated precisely through recurrent perturbations, transfer to fresh medium, dilution, abrupt renewal of carbon resources, changing oxygen availability and repeated recovery of the grain. Such disturbances are not exceptional events imposed on the system; they constitute its normal ecological cycle.

Our experiments do not demonstrate antifragility in the strict sense. To do so would require showing that controlled perturbations improve subsequent community performance, stability or adaptive capacity relative to an unperturbed state. Nevertheless, the concept provides a useful hypothesis for future experiments. Repeated perturbations could be imposed experimentally in amplitude and frequency while measuring fermentation kinetics, transition timing, grain growth and community recovery. One could then distinguish a merely robust consortium from one in which previous disturbance changes the response to subsequent perturbation. From this perspective, water kefir offers an unusually tractable model in which to investigate how ecological memory, metabolic plasticity and spatial organization combine to generate persistent community function. The consortium is sufficiently complex to display emergent behaviour, yet sufficiently constrained to be reconstructed genomically and followed over an entire fermentation cycle. Its repeated propagation furthermore makes it possible to study not only how a microbial community responds to its environment, but how its history influences that response.

Overall, our results support a view of water kefir fermentation as a coupled ecological and metabolic trajectory rather than as a simple succession of independently acting microbial groups. In the MCF consortium, the dominant dynamical feature is a reproducible reorganization of extracellular carbon metabolism around 20 h, occurring while the grain-associated community remains comparatively stable and the liquid-phase community undergoes substantial redistribution. Future work should combine finer temporal sampling, absolute population measurements, isotope-resolved carbon fluxes and controlled perturbations of individual community members to determine which interactions generate this transition and whether comparable metabolic reorganizations are conserved across independent water kefir consortia.

## 5 Acknowledgements

The authors thank Marie-Claire Frederic for providing MCF grains and her invaluable sharing of empirical experience with kefir growing over the years. We also thank Yves Barral for his suggestions to look into the GAR phenotype in the kefir consortium. Finally, we would like to thank two master students with whom we initiated this project several years ago; Nehama Edery and Anne-Claire Verschueren. The project was funded with core funds from INSERM, CNRS, Sorbonne Université and the Museum National d’Histoire Naturelle.

